# A non-invasive method to sample immune cells in the lower female genital tract using menstrual discs

**DOI:** 10.1101/2023.11.16.567469

**Authors:** M. Quinn Peters, Eva Domenjo-Vila, Marc Carlson, Blair Armistead, Paul T. Edlefsen, Melanie Gasper, Smritee Dabee, Christopher Whidbey, Heather B. Jaspan, Martin Prlic, Whitney E. Harrington

## Abstract

T cells in the human female genital tract (FGT)^2^ are key mediators of susceptibility to and protection from infection, including HIV and other sexually transmitted infections. There is a critical need for increased understanding of the distribution and activation of T cell populations in the FGT, but current sampling methods require a healthcare provider and are expensive, limiting the ability to study these populations longitudinally. To address these challenges, we have developed a method to sample immune cells from the FGT utilizing disposable menstrual discs which are non-invasive, self-applied, and low-cost. To demonstrate reproducibility, we sampled the cervicovaginal fluid (CVF)^3^ of healthy, reproductive-aged individuals using menstrual discs over three sequential days. CVF was processed for cervicovaginal cells, and high parameter flow cytometry was used to characterize immune populations. We identified large numbers of live, CD45+ leukocytes, as well as distinct populations of T cells and B cells. Within the T cell compartment, activation and suppression status of T cell subsets were consistent with previous studies of the FGT utilizing current approaches, including identification of both tissue resident and migratory populations. In addition, the T cell population structure was highly conserved across days within individuals but divergent across individuals. Our approach to sample immune cells in the FGT with menstrual discs will decrease barriers to participation and empower longitudinal sampling in future research studies.

## INTRODUCTION

In the female genital tract (FGT), the cervix and vagina are the primary sites of exposure to HIV and other sexually transmitted infections (1, 2). Local T cells at these sites are key mediators of both susceptibility to and protection from infection (3). For example, non-specific inflammation may lead to T cell activation and increased risk of HIV acquisition (2, 4–7). In contrast, robust Th1 responses are needed for clearance of pathogens such as *Chlamydia trachomatis* (8). In addition, T cells patrolling the FGT must tolerate resident microbes and other foreign antigens such as those from semen (5, 9–11). Prior phenotypic and transcriptional profiling of mucosal T cell populations has demonstrated that they are highly adapted to their site of residence and distinct from those in circulation, emphasizing the critical need to sample these populations directly from the site of interest (12–14). For example, limited work on the FGT describes an association between cervical T cell populations, menstrual cycle, and microbial community state; however, these studies have been restricted to small sample sizes (15–17). Despite the intriguing associations between the FGT and these biological variables, it has been challenging to comprehensively characterize their complex interplay in diverse populations due to the difficulty in sampling the FGT longitudinally.

Common methods for sampling immune cell populations in the FGT include biopsy, cytobrush, and cervicovaginal lavage (CVL)^4^ (18, 19). Tissue collected either from biopsy or discarded surgical tissue is considered the “gold standard” in the field for obtaining large cell yields and allows for precise sampling of anatomical sites (18, 19). This approach has identified two key T cell populations in the FGT: tissue resident memory (T_RM_)^5^ cells and migratory cells, which circulate between the tissue, regional lymph nodes, and peripheral blood (17, 20, 21). However, these important studies were restricted to small sample sizes, and most were conducted at a single time point. Cytobrushes have been used to elucidate endocervical CD4+ and CD8+ T cell populations including the impact of HIV (22–26), but the collection site is anatomically distinct from ectocervical biopsy and may preferentially sample monocytes over lymphocytes (18, 27, 28). In addition, both biopsy and cytobrush may disrupt the mucosal barrier and temporarily increase susceptibility to or transmission of infection, including HIV (29). CVL, in contrast, collects T cells present in the cervical fornixes and vaginal lumen (i.e., “luminal” T cells), and does not disrupt the mucosal barrier of the FGT. However, CVL yields substantially fewer cells than cytobrush and tissue biopsy sampling (18). Finally, each of these methods requires implementation by a health care provider in a clinical setting with a speculum exam, which can be both uncomfortable and expensive, limiting number of participants and time points for longitudinal sampling.

Here, we present a non-invasive approach to sample luminal T cells from the FGT, utilizing self-inserted menstrual discs to collect cervicovaginal fluid (CVF). Menstrual discs are a disposable personal hygiene product designed to collect menstrual fluid. In the research setting, they have been used for the study of cells present in menstrual blood (30, 31). In addition, they can be used to collect CVF in individuals while not menstruating (32–34) (**Figure 1A**, **B**). For example, menstrual discs have been used previously for the study of cytokines and other soluble markers in CVF (34–36). Given the position of the disc, menstrual discs are expected to collect both cervical secretions and fluid pooled in the vaginal fornixes. We have found that this approach yields high numbers of immune cells including T cells and captures both tissue resident and migratory populations with reproducible distributions and phenotypic features within individuals. This inexpensive, non-invasive approach does not require a clinical setting and thus reduces barriers associated with sequential sampling and empowers a larger and more diverse population of individuals to participate in future studies of FGT immunity.

**Figure 1.**
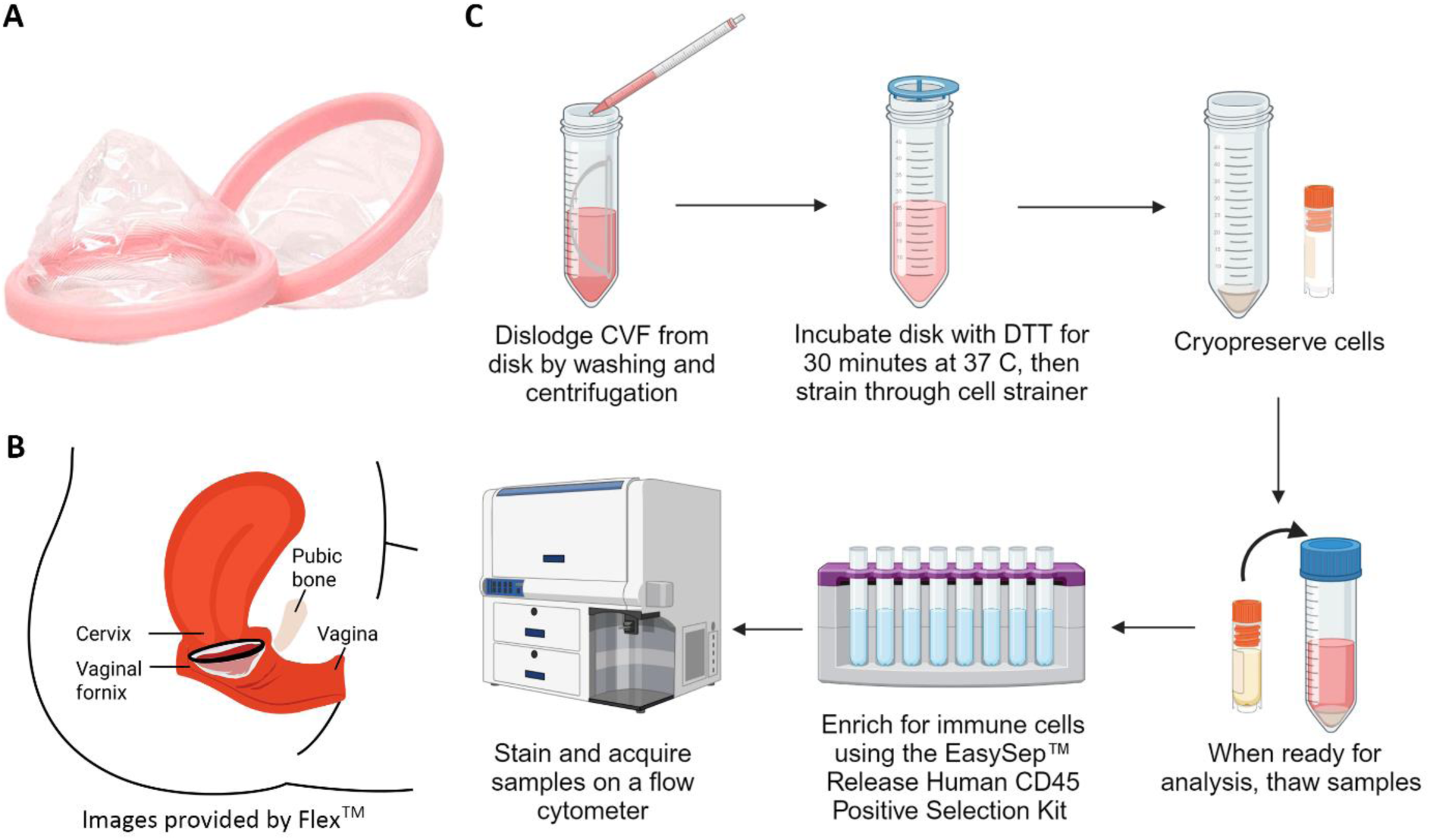
Menstrual disc sample collection and processing overview. **(A)** Softdisc® with firm plastic ring and soft plastic cup used to collect CVF. **(B)** Placement of the menstrual disc across the cervix and positioned to collect CVF. Images in panel A and B are subject to Copyright ©2023 The Flex Company, www.flexfits.com. All rights reserved. SOFTDISC is a registered trademark of The Flex Company. **(C)** Overview of method for processing and analyzing CVF for immune cells. Figure created with BioRender.com.

## MATERIALS AND METHODS

### Cohort

All participants were enrolled into the Center for Global Infectious Disease Research Biorepository, approved by the Seattle Children’s Research Institute Institutional Review Board (STUDY00002048). Participants self-reported as healthy, not pregnant, reproductive age, weighing > 110 pounds, and none had an intrauterine device or a known hormonal disorder. To validate the reproducibility of our protocol, five participants were invited to donate CVF samples on three sequential days, 7-11 days from last menstrual period (LMP) during the follicular phase, as well as one blood sample. One participant (participant 3) did not menstruate due to oral contraceptive use, so timing was not relative to LMP. Information on acute and chronic medical conditions and medication usage was also collected.

### Sample Collection

Whole blood was drawn into EDTA vacutainer tubes (BD Biosciences). Blood was processed for PBMCs as previously described (37). For CVF collection, Softdisc® menstrual discs were self-inserted by participants and worn for up to 4 hours. After self-retrieval, menstrual discs were placed in a 50 mL conical tube and returned to the study team. CVF was stored at 4°C until processing.

### Protocol for Processing of CVF From Menstrual Discs

Discs were initially processed in 50 mL conical tubes returned by participants (**Figure 1C**). Discs were submerged in 10 mL of complete RPMI (RPMI with L-glutamine, 10% fetal bovine serum (FBS), 100 U/ml penicillin, 100 µg/ml streptomycin) and rinsed 5 times in the medium using a serological pipet to dislodge CVF. The tube containing the disc was centrifuged at 250 x *g* for 7 minutes at 4°C. The menstrual disc was then removed from the tube using sterile forceps and remaining CVF was transferred from the disc to the tube by pipet. The catch basin of the menstrual disc was rinsed with 1 mL of sterile PBS with 2% FBS (FACS buffer), which was transferred to the tube, and the disc was discarded. Mucins were degraded by incubating CVF with 1 mM DTT for 30 minutes in a 37°C incubator with 5% CO_2_, as described (38). The solution was then filtered through a 100 µm nylon cell strainer (VWR) and centrifuged at 400 x *g* for 10 minutes at 4°C and the supernatant was discarded. The resulting cell pellet of cervicovaginal cells (CVC)^6^ was washed twice with 40 mL sterile PBS with 2% FBS, and cells were enumerated using a C-Chip hemocytometer (INCYTO). For cryopreservation, the cell pellet was resuspended in 1 mL freezing medium (50% FBS, 40% RPMI with L-glutamine, 10% DMSO (Millipore Sigma)), transferred to a cryovial, placed in a 1°C cryogenic freezing container at -80°C overnight, and then transferred to liquid nitrogen.

Prior to flow cytometric analysis, CVC and PBMC were thawed in a 37°C water bath, transferred to pre-warmed thaw medium (RPMI with L-glutamine, 20% FBS), and centrifuged at 400 x *g* for 5 minutes. Cell pellets were resuspended in 2 mL cRPMI and enumerated. CVF was enriched for CD45+ cells using the EasySep™ Release Human CD45 Positive Selection Kit (Stem Cell Technologies) according to manufacturer protocol. The resulting CD45 depleted fraction underwent a second enrichment to recover any remaining CD45+ cells, and the two CD45-enriched fractions were combined.

### Cell staining and Flow Cytometry Acquisition

All flow cytometry was performed at Fred Hutchison Cancer Center utilizing a 28-color panel focused on T cell phenotyping (**Supplementary Table 1**) (39). CVC and PBMC were incubated with Fc-blocking reagent (BioLegend) and fixable UV Blue Live/Dead reagent (ThermoFisher) in PBS for 15 minutes at room temperature. Cells were then stained with 50 µL of extracellular antibody master mix in Brilliant Stain Buffer (BD Bioscience) for 20 minutes at room temperature. Stained cells were washed and resuspended in FACS buffer. For intracellular and intranuclear staining, the cells were fixed with freshly prepared fixation buffer (ThermoFisher) for 30 minutes at room temperature, washed, and then stained with 50 µL of intracellular antibody master mix in FACS buffer for 20 minutes at room temperature. Cells were then washed and resuspended in FACS buffer and kept at 4°C protected from light until analysis. Single-stained compensation controls were prepared for each experiment using antibody capture beads (BD Biosciences) diluted in FACS buffer and amine reactive beads (ArC™ Amine Reactive Compensation Bead Kit, Themo Fisher). Data was acquired using a FACSymphony A5 (BD Biosciences) and FACSDiva acquisition software (BD Biosciences). All samples were run in the same experiment to eliminate batch effects.

### Flow Cytometry Analysis

Flow cytometry data were analyzed using FlowJo v10.8 software (FlowJo, LLC). The pre-determined gating scheme was applied uniformly across all samples. Initial panel development included fluorescence minus one to set gates (39). A technical replicate sample that had been validated in prior experiments with the same panel and instrument was included to inform gating. Briefly, paired CVC and PBMC were gated on CD45+ and viability, then CD3+ and CD19+ populations (**Figure 2A**). After gating out MAIT and γδ-T cells, CD3+ T cells were gated into CD4+ and CD8+ T cell populations. Within the CD4+ T cell population, regulatory T cells (Tregs) were designated as CD25+/CD127-, and the remaining cells were designated as conventional CD4+ cells. CD8+, CD4+, and Treg populations were then gated for memory, tissue residency, activation, and suppression markers. Phenotypic analysis was restricted to samples where the parent population of interest contained >=50 T cells. Cell count, frequency, and mean fluorescence intensity (MFI)^7^ were determined in FlowJo.

**Figure 2.**
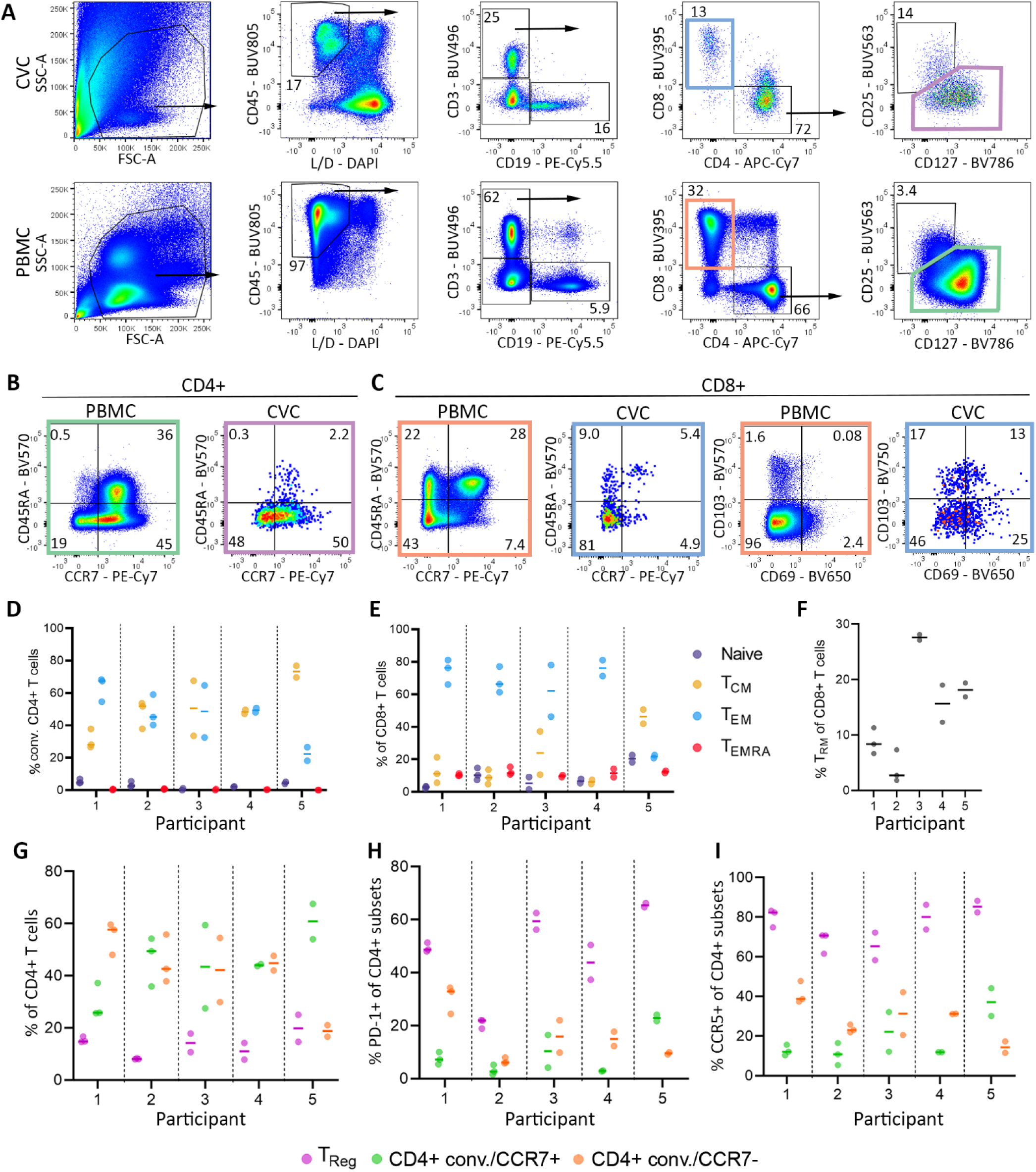
Overview of CVC T cell populations. Data from five individuals sampled across three days. Results for samples with greater than 50 cells in the parent population for each analysis are presented. (**A**) Representative gates for identification of T cells and other immune subpopulations in CVC (top) and PBMC (bottom) samples. γδ-T cells and MAIT cells were excluded from the T cell population prior to gating into CD4+ and CD8+ subpopulations. (**B**) After gating out Tregs (CD25^high^, CD127^low^), conventional CD4+ T cells were further gated by CCR7 and CD45RA to define effector memory (T_EM_; CD45RA-CCR7-), central memory (T_CM_; CD45RA-; CCR7+), terminally differentiated effector memory (T_EMRA_; CD45RA+, CCR7-), and naïve-like (CD45RA+; CCR7+) populations for PBMC (left) and CVC (right) samples. (**C**) CD8+ T cells were also gated by CCR7 and CD45RA to determine memory subtypes, as well as by CD69 and CD103 to determine tissue residency (CD69+ CD103+). (**D**) Frequency of conventional CVC CD4+ memory subtypes across samples. (**E**) Frequency of CVC CD8+ memory subtypes across samples. (**F**) Frequency of CVC CD8+ T_RM_ across samples. (**G**) Frequencies of CVC Tregs, conventional CCR7+, and conventional CCR7-cells of all CD4+ T cells. (**H**) PD-1 expression on CVC CD4+ T cell subsets. (**I**) CCR5 expression on CVC CD4+ T cell subsets.

Integrated high-parameter data analysis was restricted to CVC samples containing >=100 T cells. Comparison of global T cell population structure across samples was first conducted in R v4.3.2 using Principal Component Analysis (PCA)^8^ computed from MFI of phenotypic markers relevant to viable conventional T cells (i.e., live/dead, CD45, CD19, MR1-tet, and γδ-TCR were excluded). PCA plots were visualized using ggplot2 (40). Initial comparison included both CVC and PBMC, where only memory T cell populations were considered (i.e., naïve T cells excluded), as well as subsequent analysis considering only CVC. Comparison of local T cell population structure across all samples was performed using FlowSOM and ConsensusClusterPlus in the integrated CATALYST R package (41). T cells across CVC samples were clustered using median fluorescent intensity of the T cell-directed markers used in PCA with a maximum metacluster number of 20. The number of clusters examined for subsequent analysis (k=13) was determined by cluster stability measure when the relative change in area under a Cumulative Distribution Function curve approached zero. The aggregated marker expression within T cell clusters was scaled from 0 to 1 and visualized with a heatmap; hierarchical clustering is shown with dendrograms representing Euclidean distance and plotted with average linkage. Relative abundances of the clusters in each CVC sample were plotted to demonstrate reproducibility in population structure within individuals across samples. Fluorescent intensity data for each cell was dimensionally reduced and visualized using uniform manifold approximation and projection (UMAP)^9^ without the default CATALYST data transformation. The resulting UMAP was colored by FlowSOM assigned clusters.

### Statistical Analysis

Cell count data (e.g., number of CD3+ T cells recovered) are presented as medians. Frequencies of parent populations (e.g., % Tregs of CD4+ T cells) are presented as means. To compare frequency of populations between CVC and PBMC, a mean frequency of the CVC population across all sampled days was computed. The mean frequency of the population in the CVC was then compared with the frequency in paired PBMC. Given the small sample size and paired data structure, p values were computed with a Wilcoxon signed rank test. Significance was defined as a p value less than or equal to 0.05. Analysis was conducted in Stata 14.2.

## RESULTS

### Overview of Immune Cells Recovered from CVF

Total number of viable CD45+ cells recovered from CVF samples varied both between and within individuals (**Table 1**). The median number of viable CD45+ cells recovered from CVF was 6,235. Within the CD45+ cell population, CD3+ T cells were the largest subpopulation (median = 1,182 cells), but there was also high recovery of CD19+ B cells (median = 1,072) (**Table 1**). Of CD3+ T cells, frequencies ranged from 37-87% for CD4+ T cells and 7-55% for CD8+ T cells. The majority of CVC T cells were antigen-experienced (mean: 97% of conventional CD4+ cells and 91% of CD8+ cells), including central memory (CCR7+ CD45RA-), effector memory (CCR7-CD45RA-), and terminally differentiated effector CD45RA+ populations (CCR7-, CD45RA+) **(Figure 2B-E)**. The CD8+ compartment of CVC contained both T_RM_ (CD69+ CD103+) (**Figure 2F**) and migratory populations, and CD8+ T_RM_ were significantly enriched in CVMC relative to PBMC (mean: 15% vs. 0.03%, p=0.04). In addition to CCR7+ and CCR7-conventional populations, CVC CD4+ T cells also contained a Treg population (**Figure 2G**), the frequency of which was enriched compared to PBMC (mean: 14% vs 4%, p=0.04). Similar to conventional CD4+ T cells, 98% of Tregs were antigen experienced. Within CVC CD4 T cells, the expression of PD-1 and CCR5 was highest on Tregs, relative to conventional CD4+ T cells, and was consistent across sampling days within individuals (**Figure 2 H-I**).

**Table 1.** Counts of immune cell populations across CVF samples

| Participant | 1 |  |  | 2 |  |  | 3 |  |  | 4 |  |  | 5 |  |  |  |
| --- | --- | --- | --- | --- | --- | --- | --- | --- | --- | --- | --- | --- | --- | --- | --- | --- |
| Sample | 1 | 2 | 3 | 1 | 2 | 3 | 1 | 2 | 3 | 1 | 2 | 3 | 1 | 2 | 3 | Median |
| CD45+ | 14,772 | 11,157 | 57,703 | 4,533 | 13,401 | 20,587 | 1,225 | 2,036 | 11,860 | 6,233 | 212 | 31,479 | 2,885 | 965 | 2,762 | 6,233 |
| CD3+ | 3,174 | 1,182 | 11,554 | 1,052 | 1,966 | 3,512 | 38 | 392 | 3,897 | 1,808 | 12 | 7,823 | 521 | 56 | 384 | 1,182 |
| CD4+ | 2,037 | 938 | 7,484 | 518 | 1,197 | 1,460 | 14 | 223 | 2,037 | 1,567 | 5 | 5,449 | 350 | 34 | 271 | 938 |
| CD8+ | 902 | 157 | 3,404 | 191 | 331 | 659 | 21 | 118 | 1,387 | 163 | 2 | 968 | 77 | 4 | 31 | 163 |
| CD19+ | 2,166 | 727 | 3,638 | 1,072 | 2,070 | 2,599 | 51 | 456 | 1,220 | 1,548 | 8 | 4,690 | 170 | 25 | 201 | 1,072 |
| CD3/CD19-<br>HLA-DR+ | 5,539 | 4,082 | 12,849 | 605 | 3,052 | 3,726 | 482 | 392 | 3,019 | 611 | 32 | 3,251 | 775 | 64 | 452 | 775 |

### Activation and Suppression Phenotype of CVC T cells

We next compared the activation and suppression phenotype of specific FGT T cell populations of interest reported in prior studies, specifically CD4+ T regs and memory CD8+ T cells (20, 42). Relative to PBMC, a higher frequency of CVC CD4+ Tregs expressed the activation marker CCR5 (mean: 77% vs. 39%, p=0.04) and the suppressive markers ICOS (mean: 54% vs 21%, p=0.04), PD-1 (mean: 47% vs 5%, p=0.04), LAG-3 (mean: 30% vs. 4%, p=0.04), CTLA-4 (mean: 38% vs. 7%, p=0.04), and TIM-3 (mean: 42% vs. 4%, p=0.04) (**Figure 3A**, **B**). The Treg compartment also contained populations of PD-1+ TIM3+ cells (10 to 45% within samples), a highly suppressive phenotype observed in tissue Treg populations and rarely observed in the periphery (43). Within CVC memory CD8+ T cells there was a higher frequency of cells expressing CCR5 (mean: 85% vs. 25%, p=0.04), HLA-DR (activation) (mean: 40% vs 11%, p=0.04), and PD-1 (mean: 34% vs. 7%, p=0.04) relative to PBMC (**Figure 3C**, **D**). Although the frequencies of memory CD8+ T cells expressing CD38 (mean: 23% vs 11%, p=0.08) and granzyme B (mean: 32% vs. 25%, p=0.7) was higher in CVC versus PBMC, these differences did not reach significance. Together these observations indicate that menstrual disc sampling yielded key populations of CD4 and CD8 T cells with similar levels of activation and suppression as previously reported from current sampling methods.

**Figure 3.**
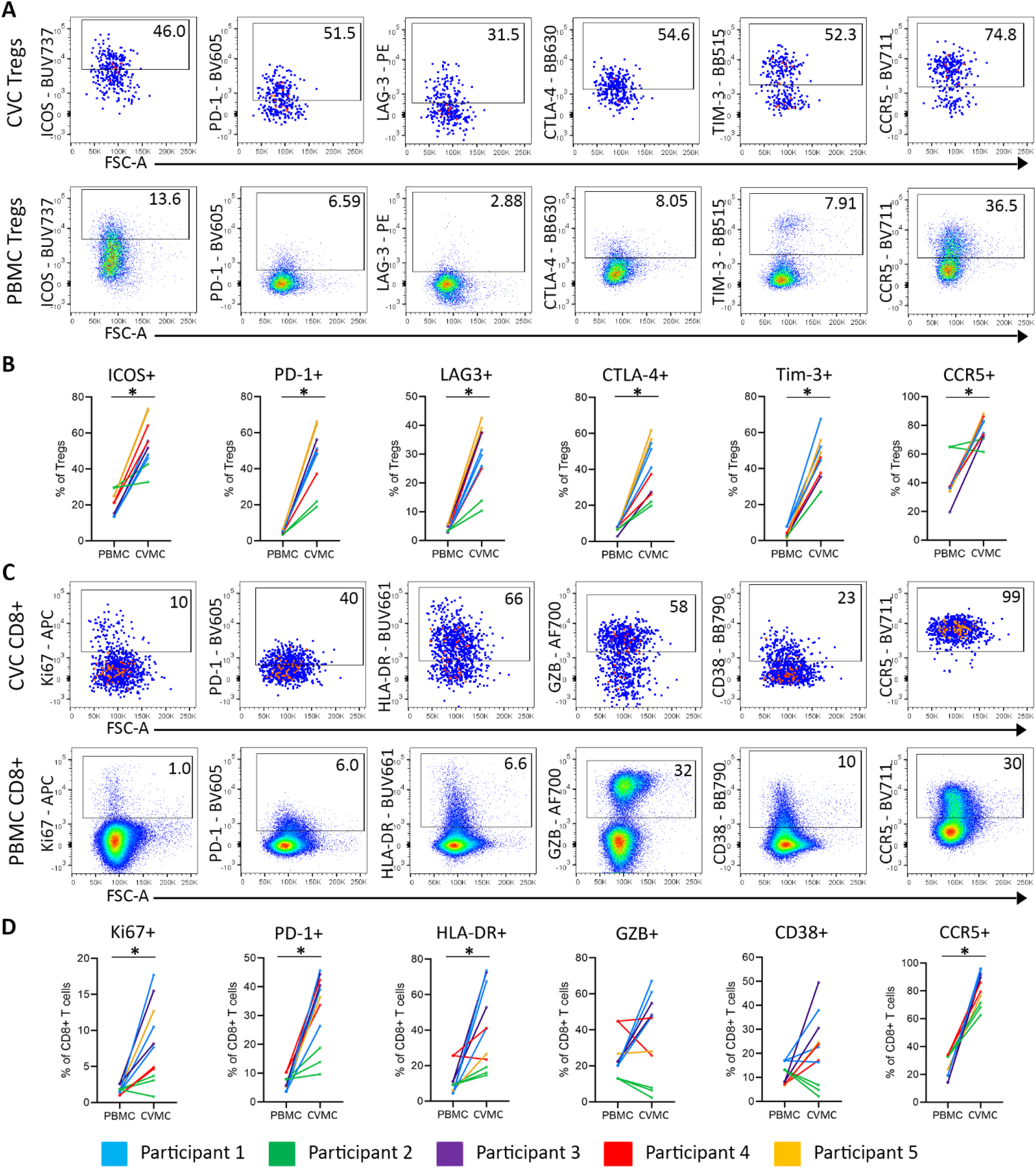
Characterization of key CVC T cell populations. Data from five individuals sampled across three (CVC) and one (PBMC) day. Results for samples with greater than 50 cells in the parent population for each analysis are presented. The mean CVC frequency of each population of interest across sampled days was compared to the PBMC frequency using a Wilcoxon signed rank test. Star indicates p value less than or equal to 0.05. **(A)** Representative flow plots of activation and suppression marker expression from paired CVC and PBMC CD4+ Treg populations. **(B)** Summary plots of frequencies of activation and suppression markers between paired CVC and PBMC CD4+ Treg populations. **(C)** Representative flow plots of activation and suppression marker expression from paired CVC and PBMC memory CD8+ T cells. **(D)** Summary plots of frequencies of activation and suppression markers between paired CVC and PBMC memory CD8+ T cells.

### Reproducibility of T cell Population Structure

To quantify the reproducibility of global CVC T cell population structure across sequential timepoints within individuals, we compared memory T cell phenotypes across consecutive days from the FGT, as well as paired PBMC from one timepoint, using PCA. When comparing CVC and PBMC memory T cell populations, samples clustered clearly by sample type (**Figure 4A**). Within CVC CD3+ and CD4+ T cell populations, by comparison, samples clustered by individual (**Figure 4B**, **C**). There was no clear relationship between day of sampling and clustering pattern for any analysis. Within CD3+ populations, the largest contributors to PC^10^1 as determined by loading scores were CD4, activation markers (CD28 and CD38), and suppressive markers (PD-1, CTLA-4, ICOS, and TIM3); the largest contributors to PC2 were memory markers (CD45RA, CCR7, and CD27) and the proliferation marker KI67. Within the CD4+ population, the largest contributors to PC1 were activation markers (CCR5, CD69, CD38, and CD28) and suppressive markers (PD-1, CTLA-4, and TIM3); the largest contributors to PC2 were also activation (HLA-DR, CD137, CD25) and suppressive markers (PD-1, LAG3). Together these observations emphasize that individuals have unique patterns of memory, activation, and suppression markers on CVC T cells that are highly conserved across days.

**Figure 4.**
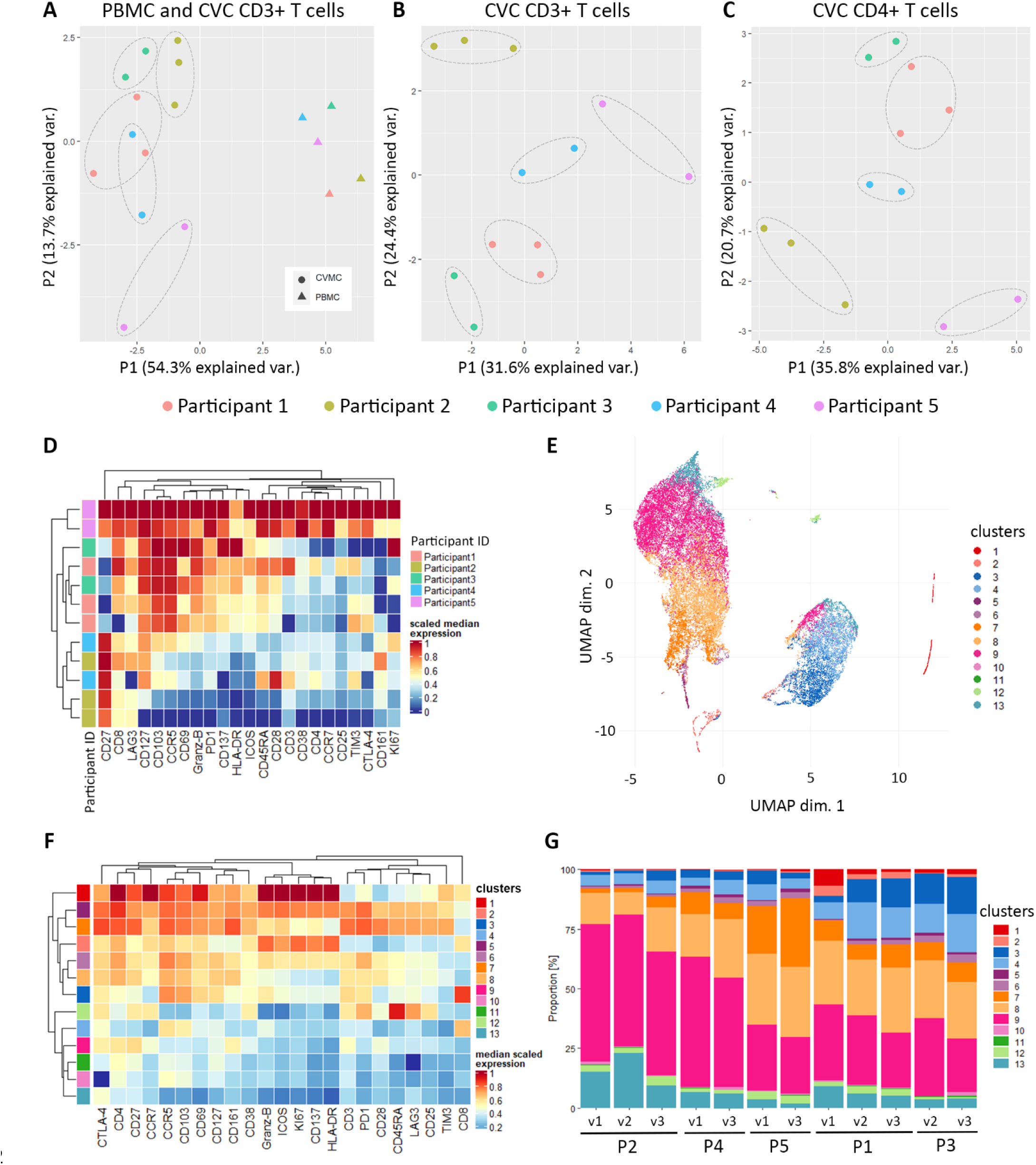
CVC T cell populations are reproducible within individuals but vary across individuals. Data from five individuals sampled across three (CVC) and one (PBMC) day. Results for samples with greater than 100 cells in the parent population for each analysis are presented. **(A)** PCA utilizing high parameter flow data to characterize paired CVC and PBMC CD3+ memory T cell populations. **(B)** PCA characterizing CVC CD3+ memory T cell populations. **(C)** PCA characterizing CVC CD4+ memory T cell populations. **(D)** Unsupervised hierarchical heat map characterizing the median expression of CVC T cell markers across individuals and samples. **(E)** Dimensional reduction of all markers relevant to T cells was performed and is represented as a UMAP including all CD3+ T cells across CVC samples. **(F)** Heat map characterizing the median expression of T cell markers in each identified cluster. **(G)** Proportions of cell clusters composing each sample, arranged by participant and visit.

We next examined local CVC T cell population structure across participants using the CATALYST tool. Consistent with our PCA analysis, initial analysis looking at marker expression demonstrated two groups of participants separated by degree of T cell activation (e.g., CCR5 expression was high in participant 1, 3, and 5 and low in participant 2 and 4) (**Figure 4D**). We next clustered T cells across all individuals to understand shared population structure. Across all participants, luminal T cells were comprised of two large populations (CD4+ and CD8+) and a few minor populations, which could be subdivided into 13 phenotypic clusters (**Figure 4E**, **F**). Clusters 1, 5, and 7 had high expression of CD4 and high expression of activation markers including CCR5, CD69, and CD137 and the tissue residency marker CD103. In particular, cluster 1 had exceptionally high activation and was distinct from the other CD4 clusters. Clusters 6, 8, and 9 were also CD4 predominant, but with lower degrees of activation and variable expression of CD45RA. Cluster 10 and 11 had CD4 expression, but showed very low activation, with cluster 10 displaying very high CTLA-4 expression and cluster 11 very high LAG-3 expression. Cluster 2 contained both CD4 and CD8 populations, with moderate activation and no expression of suppression markers. Clusters 3 and 4 were CD8 predominant, and both expressed CD103 consistent with T_RM_ populations. Cluster 12 and 13 were quite distinct with low expression of both CD4 and CD8. Within individuals, the proportion of each cluster was recapitulated across timepoints (**Figure 4G**). Together these observations indicate that luminal T cells from the FGT are comprised of diverse populations which vary by tissue residence, activation, and suppressive status and that the distribution of these populations within individuals is preserved across sequential days.

## DISCUSSION

The cervix is a critical site for exposure to and acquisition of HIV and other sexually transmitted infections. It is thus essential to understand the immune cell populations at this site, as well as how these populations change longitudinally and in response to variables such as hormonal fluctuations, microbial communities, hygiene practices, and sexual behaviors. Here, we describe a non-invasive method for obtaining large numbers of live CD45+ immune cells from CVF using menstrual discs. These samples are rich in T cells and an abundant source of B cells and HLA-DR+ populations, which remain viable after cryopreservation. The number of T cells we recovered was similar to cytobrush, which recovers approximately 500 to 5,000 T cells (18, 26), but lower than tissue biopsy which recovers approximately 5,000 to 10,000 T cells (18). Of note, compared with prior reports of cytobrush and biopsy (18, 44, 45), the observation of significant numbers of CD19+ B cells appears unique to our sampling approach.

To compare our FGT sampling method to those reported in the literature, we conducted in depth phenotyping of the T cell populations. We identified FGT T cell populations that closely resembled those previously identified in biopsy and surgically resected tissue, including both migratory and tissue resident populations (20, 42). Consistent with prior reports, both conventional CD4 and CD8 populations had high expression of CCR5, indicating activation, and for CD4+ cells representing an HIV target cell population (17, 20). In cytobrush samples, prior reports have found that approximately 65 to 75% of all CD4+ T cells are CCR5+ (26, 46), similar to our observation that 57% of all CD4+ T cells expressed CCR5. In addition, Tregs were enriched in CVC and showed high expression of suppressive markers, consistent with prior reports (42). Similarly, we identified CD8+ T_RM_, although the overall frequency was lower relative to those reported from biopsy (13, 20, 21). These data indicate that our sampling approach identifies key T cell subsets with phenotypic features consistent with prior approaches.

Importantly, our sampling method collects luminal T cells, similar to prior work utilizing CVL. In addition to representing the underlying tissue, data from lung suggest that luminal T cells may contain unique populations, which have discrete function via their ability to migrate between the lumen and the tissue (47–49). More recently, studies using CVL in non-human primates have identified a unique population of memory CD4+ T cells that migrate into the lumen of the FGT in a CCR5-dependent manner (17). In the context of the cervix and vagina, luminal T cells may preferentially encounter bacteria present as part of vaginal microbiota versus pathogenic bacteria and viruses, which invade the tissue. In this light, independent study of luminal T cells, not merely as a representation of tissue, may yield novel insights into the immunology of the cervix and vagina.

Intra-individual reproducibility is critical for any sampling method, so we assessed the reproducibility of our approach across three sequential days. Despite variability in absolute viable leukocyte recovery, the T cell population structure was maintained within each individual. Within individuals, we identified high reproducibility of both the global local population structure, including the distribution of CVC T cell populations, and specific phenotypic characteristics (e.g., the expression of PD-1 on conventional CD4+ T cells). In contrast, we observed significant inter-individual variability, wherein each participant had a unique T cell population structure which was reproducible across time. These observations suggests that differences observed across individuals may be driven by biological or behavioral variability. This highlights the need for further study to evaluate the role of specific factors in driving CVC T cell population structure across individuals.

Our study has a number of limitations. For example, the absolute number of leukocytes collected varied across days. This may reflect differences in cervical secretions day to day, variable positioning of the disc, or changes in activity level. Reassuringly, despite variation in total collected leukocytes, the relative population structure was preserved. Future implementation of this approach should consider the need for repeat sampling to ensure sufficient cell recovery for the intended application. Further, we did not collect sufficient T cells to perform both phenotypic and functional work and elected to first perform deep phenotyping. Future studies should address functional capacity as well as antigen specificity of collected T cells. In addition, different sampling methods will sample different anatomical zones and yield different populations. The high CD4:CD8 cell ratio of our CVC T cells is most consistent with cervical origin (18, 21, 27), but it should be noted that CVF collected via menstrual disc contains cells which are of less precise anatomical origin than endocervical cytobrush or tissue biopsy. Our flow cytometry panel was primarily targeted at T cell phenotyping, so we were not able to assess the relative lymphocyte to monocyte frequency; however, we did identify a large population of CD3-CD19-HLA-DR+ cells that fell within the expected forward and side scatter of monocytes (18). Finally, participants may be unfamiliar with the use of menstrual discs, with a risk of incorrect placement in the vagina rather than surrounding the cervix. To address this in future studies, study team members could demonstrate correct positioning of the disc prior to utilization.

We have established a method for cervicovaginal sampling that is low cost, non-invasive, and self-administered. This method can be used in longitudinal studies and does not require study participants to interact with a health care provider or a clinical setting, reducing the barrier to participation. The high population reproducibility, low cost, and ability to cryopreserve these samples with sufficient cell recovery further emphasizes the potential application of this approach in future studies, such as in large scale vaccine trials that may take place at multiple sites.

## Acknowledgements

We thank the study participants who donated blood and CVF.

## Author Contributions

Methodology: MQP, EDV, MC, PTE, MG, SD, HBJ, MP, CW, WEH

Participant enrollment: MQP, WEH

Investigation: MQP, WEH

Visualization: MQP, MC, BA, WEH

Formal Analysis: MQP, MC, PTE, SD, WEH

Funding Acquisition: MQP, BA, WEH

Supervision: MP, WEH

Writing—Draft Preparation: MQP, WEH

Writing—Review and Editing: MQP, EDV, BA, MC, PTE, MG, SD, HBJ, MP, CW, WEH

## Footnotes

1 Burroughs Wellcome Fund Career Award for Medical Scientists 1017213 National Institutes of Health K08AI135072 Institute of Translational Health Sciences, University of Washington, Early Investigator Catalyst Award BPO 75-408 University of Washington and Seattle Children’s Research Institute Seed Funds

2 FGT: Female genital tract

3 CVF: Cervical vaginal fluid

4 CVL: Cervicovaginal lavage

5 TRM: Tissue resident memory

6 CVC: Cervicovaginal cells

7 MFI: mean fluorescence intensity

8 PCA: Principal component analysis

9 UMAP: Uniform manifold approximation and projection

10 PC: Principal component

