## Supplemental Table 1 for "A non-invasive method to sample immune cells in the lower female genital tract using menstrual discs"

**Supplementary Table 1. High dimensional flow cytometry panel used for general T cell characterization.**

| | Laser | Bandpass Filter | Fluorophore | Antigen |
|---|---|---|---|---|
| 1 | 355 nm | 379/28 | BUV395 | CD8 |
| 2 | 65 mW | 450/50 | UV Blue Live/Dead | Amine Reactive |
| 3 | | 515/30 | BUV496 | CD3 |
| 4 | | 585/30 | BUV563 | CD25 |
| | | 610/20 | - | - |
| 5 | | 670/30 | BUV661 | HLA-DR |
| 6 | | 740/35 | BUV737 | ICOS |
| 7 | | 820/60 | BUV805 | CD45 |
| 8 | 406 nm | 431/28 | BV421 | MR1-Tet |
| 9 | 200 mW | 525/50 | BV480 | CD28 |
| 10 | | 586/15 | BV570 | CD45RA |
| 11 | | 610/20 | BV605 | PD1 |
| 12 | | 670/30 | BV650 | CD69 |
| 13 | | 710/50 | BV711 | CCR5 |
| 14 | | 750/30 | BV750 | CD103 |
| 15 | | 780/60 | BV786 | CD127 |
| | 488 nm | 488/10 | SSC | - |
| 16 | 200 mW | 515/20 | BB515 | TIM3 |
| 17 | | 610/20 | BB630 | CTLA-4 |
| 18 | | 670/30 | BB660 | CD27 |
| 19 | | 710/50 | BB700 | CD161 |
| | | 750/30 | - | - |
| 20 | | 780/60 | BB790 | CD38 |
| 21 | 552 nm | 586/15 | PE | CD223 |
| 22 | 150 mW | 610/20 | PE-CF594 | TCRgd |
| 23 | | 670/30 | PE-Cy5 | CD137 |
| 24 | | 710/50 | PE-Cy5.5 | CD19 |
| 25 | | 780/60 | PE-Cy7 | CCR7 |
| 26 | 628 nm | 670/30 | eF660 | KI67 |
| 27 | 200 mW | 710/50 | AF700 | Granz-B |
| 28 | | 780/60 | APC-H7 | CD4 |
